# A conserved regulatory module regulates receptor kinase signaling in immunity and development

**DOI:** 10.1101/2021.01.19.427293

**Authors:** Thomas A. DeFalco, Pauline Anne, Sean R. James, Andrew Willoughby, Oliver Johanndrees, Yasmine Genolet, Anne-Marie Pullen, Cyril Zipfel, Christian S. Hardtke, Zachary L. Nimchuk

## Abstract

Ligand recognition by cell-surface receptors underlies development and immunity in both animals and plants. Modulating receptor signaling is critical for appropriate cellular responses but the mechanisms ensuring this are poorly understood. Here, we show that signaling by plant receptors for pathogen-associated molecular patterns (PAMPs) in immunity and CLAVATA3/EMBRYO SURROUNDING REGION-RELATED peptides (CLEp) in development employ a similar regulatory module. In the absence of ligand, signaling is dampened through association with specific type-2C protein phosphatases (PP2Cs). Upon activation, PAMP and CLEp receptors phosphorylate divergent cytosolic kinases, which, in turn, phosphorylate the phosphatases, thereby promoting their release from the receptor complexes. Our work reveals a regulatory circuit shared between immune and developmental receptor signaling, which may have broader important implications for plant receptor kinase-mediated signaling in general.

## INTRODUCTION

Plants deploy receptor kinases (RKs) at the cell surface to perceive their immediate environment, and thus coordinate growth, development, reproduction, and stress responses (Hohmann et al., 2017). Leucine-rich repeat (LRR)-RKs comprise the largest RK family in plants, with over 220 members in the model plant *Arabidopsis thaliana* (hereafter Arabidopsis) (Shiu and Bleecker, 2003). Several of the best-studied LRR-RKs to date function as cell-surface immune receptors, which perceive pathogen-associated molecular patterns (PAMPs) or endogenous phytocytokines to regulate immunity. In particular, the Arabidopsis LRR-RKs FLAGELLIN SENSING 2 (FLS2) and ELONGATION FACTOR TU RECEPTOR (EFR) perceive the bacterial PAMPs flagellin (or its peptide epitope flg22) and elongation factor-Tu (or its peptide epitope elf18), respectively, to regulate pattern-triggered immunity (PTI) (Gómez-Gómez and Boller, 2000; Zipfel et al., 2004; Zipfel et al., 2006).

Receptor-like cytoplasmic kinases (RLCKs) are homologous to RKs but lack an extracellular domain, and are thought to act in downstream RK signaling (Liang and Zhou, 2018). Members of the large RLCK-VII/PBS1-LIKE (PBL) family in particular have emerged as key components of LRR-RK-mediated signaling, such as the close homologs BOTRYTIS-INDUCED KINASE1 (BIK1) and PBL1, which act downstream of several immune-related RKs, including FLS2 and EFR (Veronese et al., 2006; Lu et al., 2010; Kadota et al., 2014; Li et al., 2014), and are thus key executors of PTI.

Modulation of receptor signaling is critical to prevent inappropriate activation, and previous work has implicated several protein phosphatases in the context of LRR-RK-mediated immune signaling (Park et al., 2008; Segonzac et al., 2014; Holton et al., 2015; Couto et al., 2016), including the Arabidopsis PP2Cs POLTERGEIST-LIKE 4 and 5 (PLL4 and 5) and their rice homolog XB15, which were identified as negative regulators of EFR- and XA21-mediated PTI, respectively (Park et al., 2008; Holton et al., 2015).

## RESULTS AND DISCUSSION

### A BIK1-PLL4/5 regulatory module controls PTI activation

In keeping with a negative role in PTI, *pll4 pll5* mutants exhibited accelerated kinetics of reactive oxygen species (ROS) production in response to the PAMPs elf18 and flg22 as well as to the phytocytokine AtPep1 (Figure 1A,B), indicating that like BIK1 (Couto and Zipfel, 2016), PLL4 and 5 are common components of PTI signaling downstream of multiple LRR-RKs.

**Fig 1:**
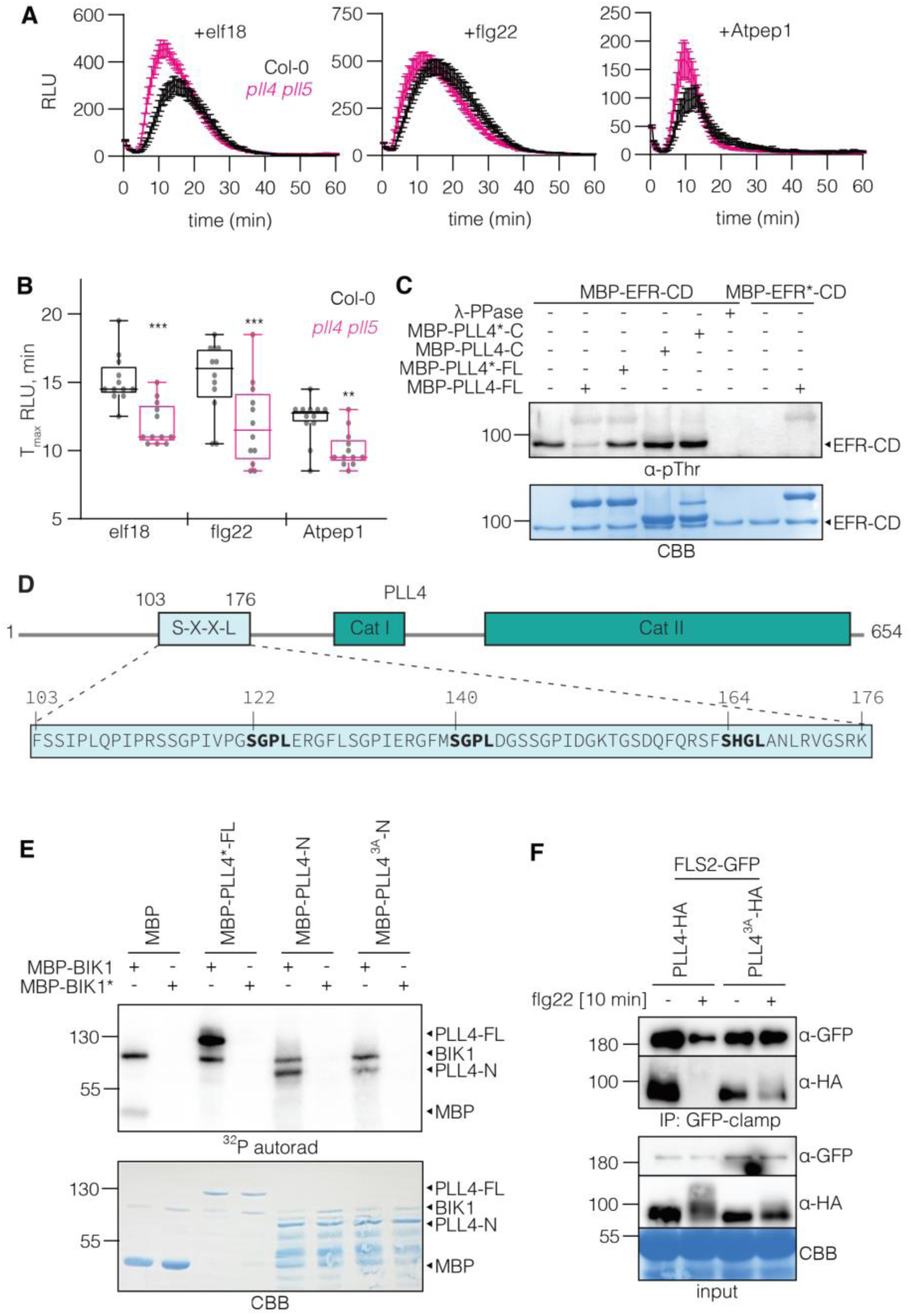
A BIK1-PLL regulatory circuit controls PTI activation. **(A-B)** PLL4 and PLL5 are involved in PAMP immune responses. ROS burst induction by elf18 (100 nM), flg22 (100 nM) or AtPep1 (1 µM) treatments on 4.5-week-old Arabidopsis leaf discs in Col-0 wildtype and *pll4 pll5* backgrounds. (A) Values correspond to the mean of 12 samples (± SE) and are expressed in relative light units (RLU). (B) Histograms represent the time to max RLU (T_max_ RLU) of 12 replicates (± SD). *** p value < 0.001, ** p value <0.01 (two-tailed T-test). **(C)** PLL4 can dephosphorylate EFR *in vitro. In vitro* phosphatase assay incubating equal amounts of MBP-tagged full length (FL) or C-terminal (C) WT (PLL4) or inactive PLL4 (PLL4*) with autophosphorylated cytosolic domain of MBP-tagged EFR (EFR-CD). The phosphorylation of MBP-EFR-CD was detected by anti-phosphothreonine western blot. **(D)** Schematic representation of PLL4 domains and details of the S-X-X-L domain of PLL4 and PLL5 homologs. Bold amino acids indicate the S-X-X-L motifs targeted for S→A mutagenesis; numbers correspond to the amino acids positions within the protein. **(E)** BIK1 phosphorylates the PLL4 N-terminus in a site-specific manner. Autoradiogram of *in vitro* kinase assay using WT BIK1 or inactive (BIK1*) BIK1 with full length (FL) or N-terminal domain (N) of WT or PLL4 phosphovariant (PLL4-N^3A^). **(F)** *In planta* flg22-triggered PLL4 dissociation from FLS2 is phosphorylation-dependent. CoIP assay of transiently expressed FLS2-GFP and HA-tagged PLL4 or PLL4^3A^ in *N. benthamiana* leaves with or without 1 μM flg22 treatment for 10 minutes. CBB: Coomassie brilliant blue.

When expressed as a maltose-binding protein (MBP)-fusion protein, wild-type (WT) but not catalytically-dead PLL4 (PLL4*, D280N/D573N) directly dephosphorylated the autophosphorylated cytosolic domain of EFR (EFR-CD) *in vitro* (Figure 1C). Interestingly, truncation of the non-catalytic N-terminus rendered PLL4 inactive in this assay, in contrast to previous work with the related phosphatase POL (Yu et al., 2003), suggesting an important role for this N-terminal region.

It was previously shown that elf18 perception induced dissociation of PLL4 and 5 from EFR *in planta* (Holton et al., 2015); however, the mechanisms mediating such dissociation remain unknown. We observed a similar flg22-induced dissociation of PLL4 and 5 from FLS2 *in planta* using transient expression in *Nicotiana benthamiana* (Figure S1), in keeping with our observation that these phosphatases also regulate FLS2 signaling (Figure 1A, B). To understand how PLL4 and 5 are themselves regulated, we interrogated public databases (Heazlewood et al., 2008; Mergner et al., 2020) for phosphosites within these proteins. Several clustered, conserved sites were identified in the N-terminus of PLL4 and 5 that conformed to a previously identified [S/T]-X-X-L motif (Figure 1D, S2), which is targeted by BIK1 and PBL1 in other substrates (Kadota et al., 2014; Thor et al., 2020). To test whether BIK1 directly phosphorylates PLL4, we expressed various fragments of PLL4 as MBP fusions and subjected them to *in vitro* kinase assays. BIK1 specifically trans-phosphorylated full-length MBP-PLL4 *in vitro*, confirming that PLL4 is a *bona fide* BIK1 substrate. This phosphorylation was constrained to the N-terminus of PLL4 (Figure 1E), and when we mutated three S-X-X-L sites (Figure 1D) to non-phosphorylatable variants (PLL4-N^3A^), phosphorylation was reduced (Figure 1E). Together, these data indicate that BIK1 specifically phosphorylates tandem sites in the PLL4 N-terminus.

To understand how PLL4 phosphorylation regulates its function, we compared ligand-induced dissociation of WT or phospho-dead mutant variants from FLS2. Treatment with flg22 induced a shift in the mobility of PLL4 indicative of phosphorylation, which was largely lost with PLL4^3A^ (Fig S3A) and flg22-induced dissociation of PLL4^3A^ from FLS2 was compromised (Figure 1F), suggesting that BIK1-directed phosphorylation regulates the dissociation of PLL4/5 from the RK complex. ROS production upon elf18 treatment was dampened by PLL4^3A^ (Figure S3B), and in keeping with a model wherein PLL4 phosphorylation triggers dissociation from receptors, phosphomimetic mutation of these sites (PLL4^3D^) impaired direct interaction with EFR-CD *in vitro* (Figure S3C).

### Isolation of the RLCK-VII-5 isoform PBL34 as a component of CLE signaling

Aside from immunity, many LRR-RKs regulate diverse growth and developmental processes, such as the CLAVATA 1 (CLV1) and BARELY ANY MERISTEM (BAM1-3) clade, which perceive endogenous peptides of the CLE family (Hazak and Hardtke, 2016; Nimchuk, 2017). CLE peptides (CLEp) are broadly conserved across land plants (Goad et al., 2017) and regulate important aspects of plant development including stem cell niche maintenance and root development (Yamaguchi et al., 2016; Fletcher, 2020). Despite the biological importance of CLEp signaling, the molecular components of such pathways are mostly unknown.

In an unbiased forward genetic screen to identify novel components of the CLEp pathway (Anne et al., 2018), two independent mutants were isolated based on their dominant insensitivity to CLE26p (Hazak et al., 2017) for which identical causative mutations were mapped to an L135F amino acid change in the RLCK-VII isoform PBL34. This dominant negative allele (*pbl34-2*) not only conferred insensitivity to CLE26p, but also to other root-active CLE peptides (Figure 2A), indicating that PBL34 is required for response to exogenous CLE peptides. Matching these observations, a *PBL34* null allele (*pbl34-3*) displayed quantitative insensitivity to the same range of root-active CLE peptides (Figure 2A). Consistent with a role downstream of CLEp perception, PBL34 and the cytosolic domain of the CLEp receptor BAM3 (BAM3-CD) were able to directly phosphorylate each other *in vitro* (Figure 2B).

**Fig 2:**
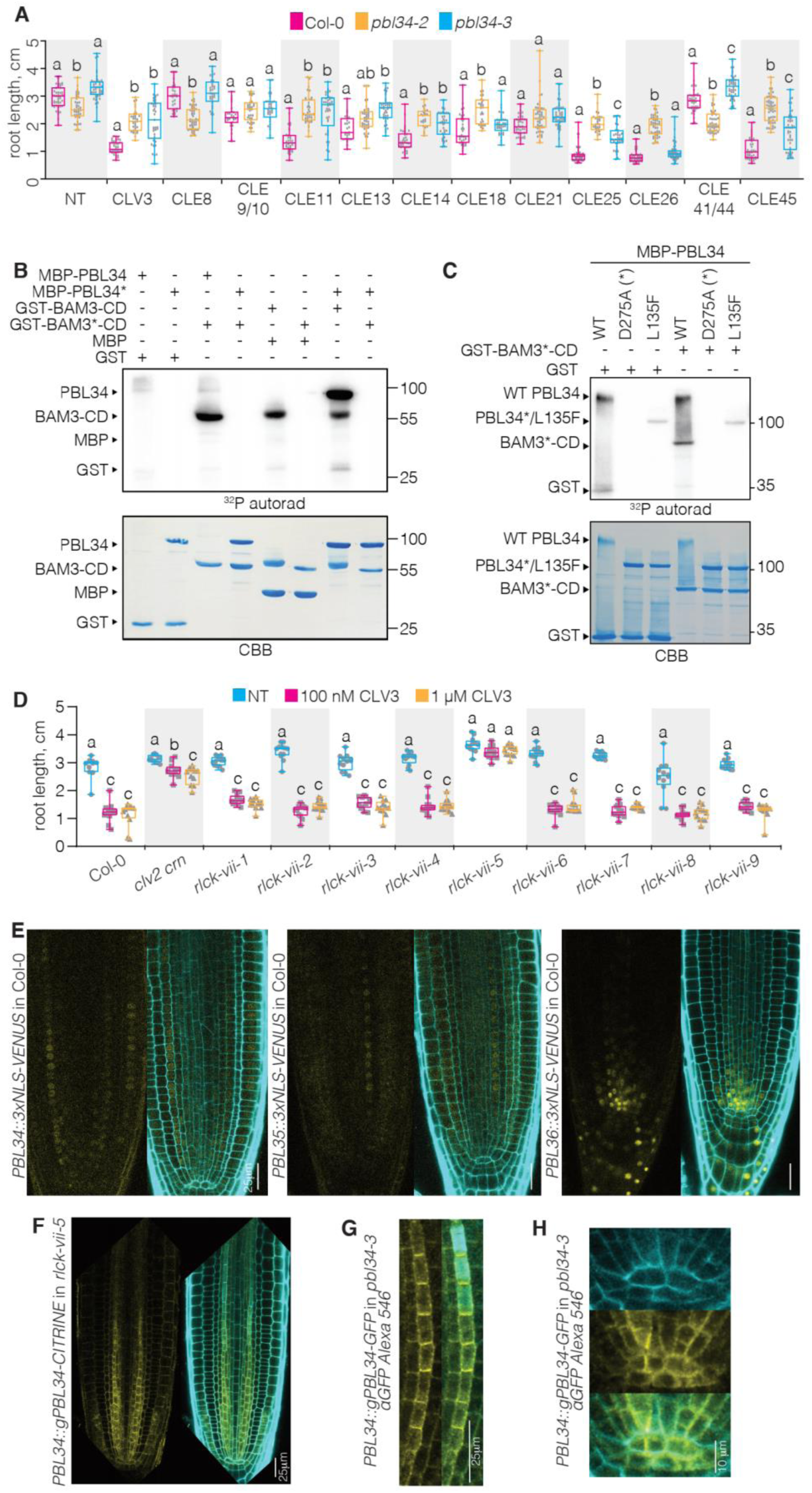
RLCK-VII-5 isoforms are required for CLE signaling. **(A)** The *pbl34-2* dominant negative allele is less sensitive to exogenous CLEp than the *pbl34-3* loss-of-function allele. 7-day-old seedlings grown on media with 50 nM of indicated CLE peptides. NT: not treated. Letters indicate significant differences within the treatments (ANOVA followed by Tukey test). n=11-50. **(B)** BAM3 and PBL34 transphosphorylate each other *in vitro*. Autoradiogram of *in vitro* kinase assay using MBP-tagged WT PBL34 or inactive PBL34 (PBL34*) and GST-tagged WT cytosolic domain (CD) of BAM3 (BAM3-CD) or inactive CD of BAM3 (BAM3*-CD). **(C)** The L135F mutation disrupts auto- and trans-phosphorylation activity of PBL34. Autoradiogram of *in vitro* kinase assay incubating equal amounts of GST-tagged BAM3 with MBP-tagged WT PBL34 or mutant forms of PBL34 (PBL34^D275A^ or PBL34^L135F^). **(D)** CLV3p responses specifically require the *RLCK-VII-5* subfamily. 7-day-old seedlings grown on media with CLV3p as indicated. NT: not treated. Letters indicate significant differences within the treatments (ANOVA followed by Tukey test). n=26-46. **(E)** *RLCK-VII-5* members are expressed in the root with partially overlapping patterns. Confocal microscopy pictures of 6-day-old seedlings carrying *PBL34::3xNLS-VENUS, PBL35::3xNLS-VENUS* and *PBL36::3xNLS*-*VENUS* constructs, respectively in Col-0 background. yellow channel: 3xNLS-VENUS; cyan: propidium iodide cell wall staining. **(F-H)** *PBL34* is expressed in the root, accumulates in the protophloem and localizes to the cytosol and the plasma membrane. Confocal microscopy images of 5-day-old seedlings expressing (G) PBL34-CITRINE fusion protein under control of the *PBL34* promoter in the *rlck-vii-5* triple mutant. Live imaging; yellow channel: CITRINE; cyan: propidium iodide cell wall staining. (H-I) Immunolocalization of PBL34-GFP protein fusion expressed under control of the *PBL34* promoter in *pbl34-3* mutant using anti-GFP primary antibody combined with Alexa 546 fluorophore (yellow channel). Cyan: calcofluor white cell wall staining. CBB: Coomassie brilliant blue.

L135 of PBL34 is highly conserved across the RLCK-VII/PBL family (Figure S4). To understand the dominant negative effect of the *pbl34-2* allele, we expressed recombinant PBL34 bearing the causative L135F mutation and tested its kinase activity *in vitro*. Both auto- and trans-phosphorylation activities of PBL34^L135F^ were severely reduced compared to WT (Fig 2C), indicating that kinase activity is essential for PBL34 function in CLE signaling.

### The RLCK-VII-5 subfamily is required for CLE signaling downstream of multiple receptors

PBL34 belongs to the RLCK-VII-5 subfamily together with PBL35 and 36 (Rao et al., 2018). Comparison of single and double mutants in these *PBL* genes revealed that each contributes quantitatively to CLEp sensitivity in root elongation assays (Figure S5A, B). Transcriptional reporters indicated expression of all three *RLCK-VII-5* isoforms in the root (Figure 2E). A loss-of-function mutant of all three isoforms (*rlck-vii-5*) led to strongly increased CLEp insensitivity (Figure 2C S5A-C), comparable or superior to the *pbl34-2* mutant (Figure S5D). A screen of all *rlck-vii* subfamily polymutants (Rao et al., 2018) revealed that this CLEp insensitivity was unique to *rlck-vii-5*, indicating that RLCK-VII-5 subfamily PBLs are specifically required for CLEp signaling (Figure 2D). The *rlck-vii-5* mutant was complemented by a PBL34-CITRINE fusion protein expressed under control of the *PBL34* promoter (Figure S5E-G), corroborating the predominant role of PBL34 within the clade. Finally, PBL34-CITRINE displayed plasma membrane association (Figure 2F-H), consistent with a role in CLEp signaling immediately downstream of CLEp receptors.

We next examined the function of RLCK-VII-5 clade kinases in specific CLEp-dependent processes controlled by the primary CLV1 and BAM1-3 LRR-RKs. Consistent with a role in primary CLEp receptor function, CLE45p inhibited protophloem differentiation in a *BAM3* (Depuydt et al., 2013) and *RLCK-VII-5* dependent manner (Figure 3A). CLE40p regulates root quiescent center (QC) stem cell maintenance (Stahl et al., 2009), where *BAM1, BAM2* (Crook et al., 2020), and *RLCK-VII-5* are expressed (Figure 2E, H). CLE40p promoted loss of quiescence and induced QC cell divisions in WT, but not in *rlck-vii-5* or *bam1 bam2* mutants (Figure 3B,C). Interestingly, while PLL4 and 5 regulate immunity, *pol* and *pll1* mutants were first identified as partial suppressors of *clv1* (Yu et al., 2000; Song and Clark, 2005), suggesting that they negatively regulate CLE signaling. Ectopic QC divisions were observed in *pol* single mutants in the absence of CLE40p (Figure 3 B,C), phenocopying CLEp treatment in WT plants.

**Fig 3:**
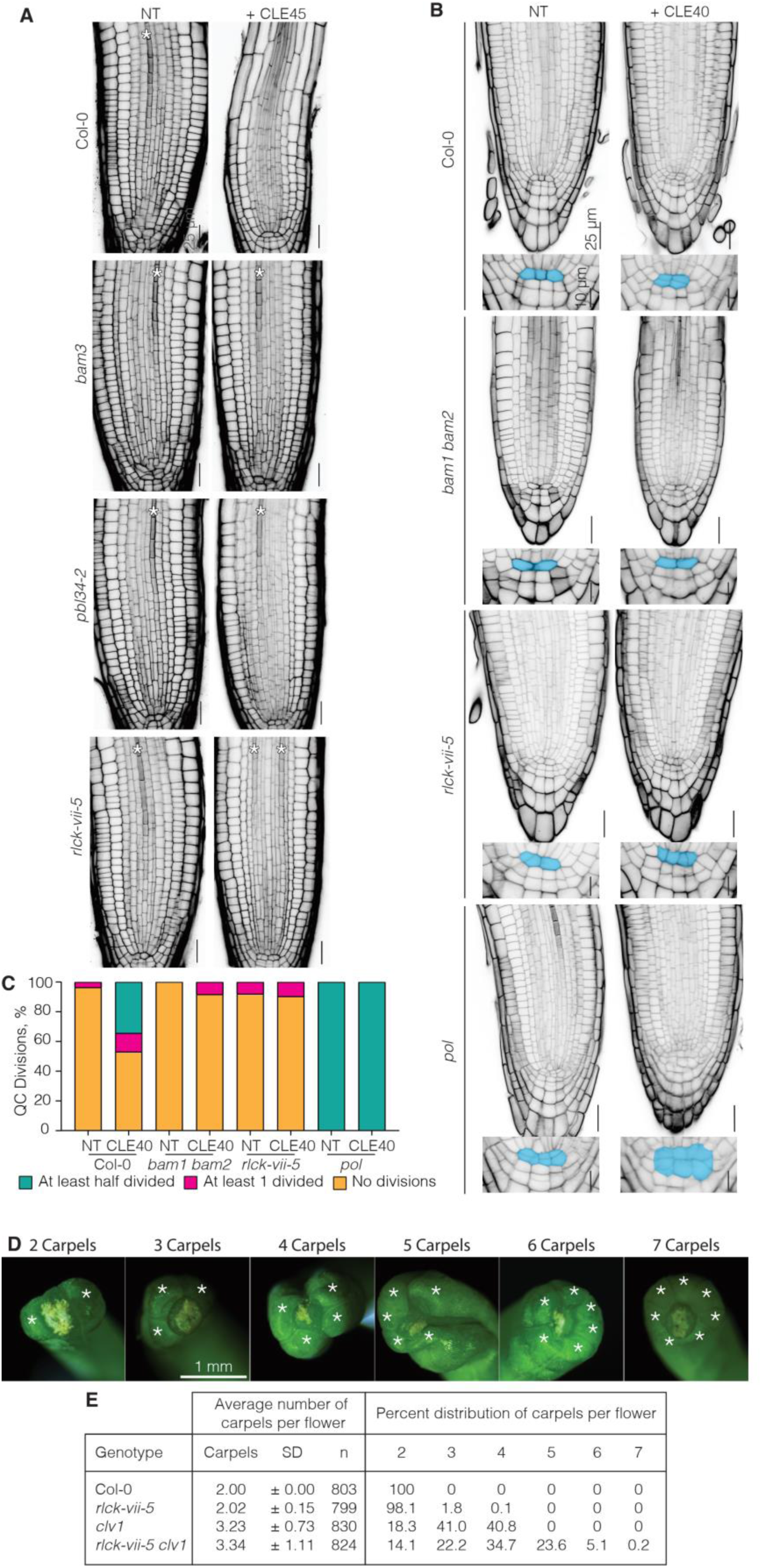
The RLCK-VII-5 subfamily functions downstream of multiple primary CLE receptors. **(A)** RLCK-VII-5 kinases control phloem differentiation through CLE45p/BAM3 signaling pathway. Confocal pictures of 5-day-old-seedlings. In *pbl34* mutants differentiated phloem files could be observed frequently, which was never observed in Col-0 wildtype. *bam3* and *rlck-vii-5* are totally CLE45p-resistant. White asterisks: phloem cell files. **(B-C)** RLCKV-VII kinases control CLE40p/BAM1/BAM2/POL mediated QC division. CLE40p triggers QC division dependent on RLCK-VII-5 kinases and BAM1 BAM2; while untreated *pol* display ectopic QC divisions mimicking CLE40p treatment (B) Representative pictures of confocal microscopy of 5-day-old seedlings treated or not with 100 nM CLE40p. black: propidium iodide cell wall staining. (C) Corresponding quantification of QC division phenotypes in control or 100 nM CLE40p treated plants. **(D-E)** *rlck-vii-5* mutants display a mild extra carpel phenotype by itself and enhances the *clv1* single mutant. (D) Representative images of carpels per flower with white asterisk indicating a carpel and (E) table representing the average number of carpels per flower (+/-standard deviation) for every flower on the primary inflorescence of 30 individual 6-week-old plants per genotype and the percent distribution of carpel number per flower.

In shoot and floral meristems, CLV3p and highly redundant CLE peptides signal through CLV1 and BAM receptors to limit stem cell proliferation (Nimchuk, 2017; Rodriguez-Leal et al., 2019). Disruption of CLV3p/CLEp signaling thereby results in increased floral organ numbers, which is reverted in *pol* or *pll1* single mutants (Song et al., 2006). Consistent with a general role of RLCK-VII-5 isoforms in CLEp perception, *rlck-vii-5* also displayed a mildly increased carpel number, which was never observed in WT plants; the *rlck-vii-5* mutant further dramatically enhanced carpel numbers in a sensitized *clv1/bam* mutant background (Figure 3D-E). Collectively, these data demonstrate that RLCK-VII-5 PBL kinases are critical for diverse CLEp-CLV1/BAM developmental outputs.

### PBL34 regulates POL via direct phosphorylation

The clustered S-X-X-L phosphosites identified in PLL4 are highly conserved across PLL isoforms (Figure S2). Given the similar roles of POL, PLL1 and PBL34 in CLEp signaling and that of PLL4, 5 and BIK1, PBL1 in PTI, we reasoned that the regulatory mechanisms could be conserved. To test this, we performed *in vitro* kinase assays with recombinant MBP-tagged POL fragments, and confirmed that PBL34 directly phosphorylates the N-terminus of POL (Figure S6A), which is lost when seven tandem putative phosphosites (Figure 4A) were mutated to non-phosphorylatable variants (POL-N^7A^) (Figure 4B). We tested the role of these phosphosites in the association between POL and CLE receptors, and observed that a phosphomimetic variant of POL (POL^7D^) was compromised in association with both BAM3 (Figure 4C) and CLV1 (Figure S6B) *in planta*, as well as showed reduced direct interaction with BAM3-CD *in vitro* (Figure S6C). In keeping with their shared role in CLEp signaling, PBL34 also specifically phosphorylated PLL1 at its N-terminus (Figure S6D).

**Figure 4:**
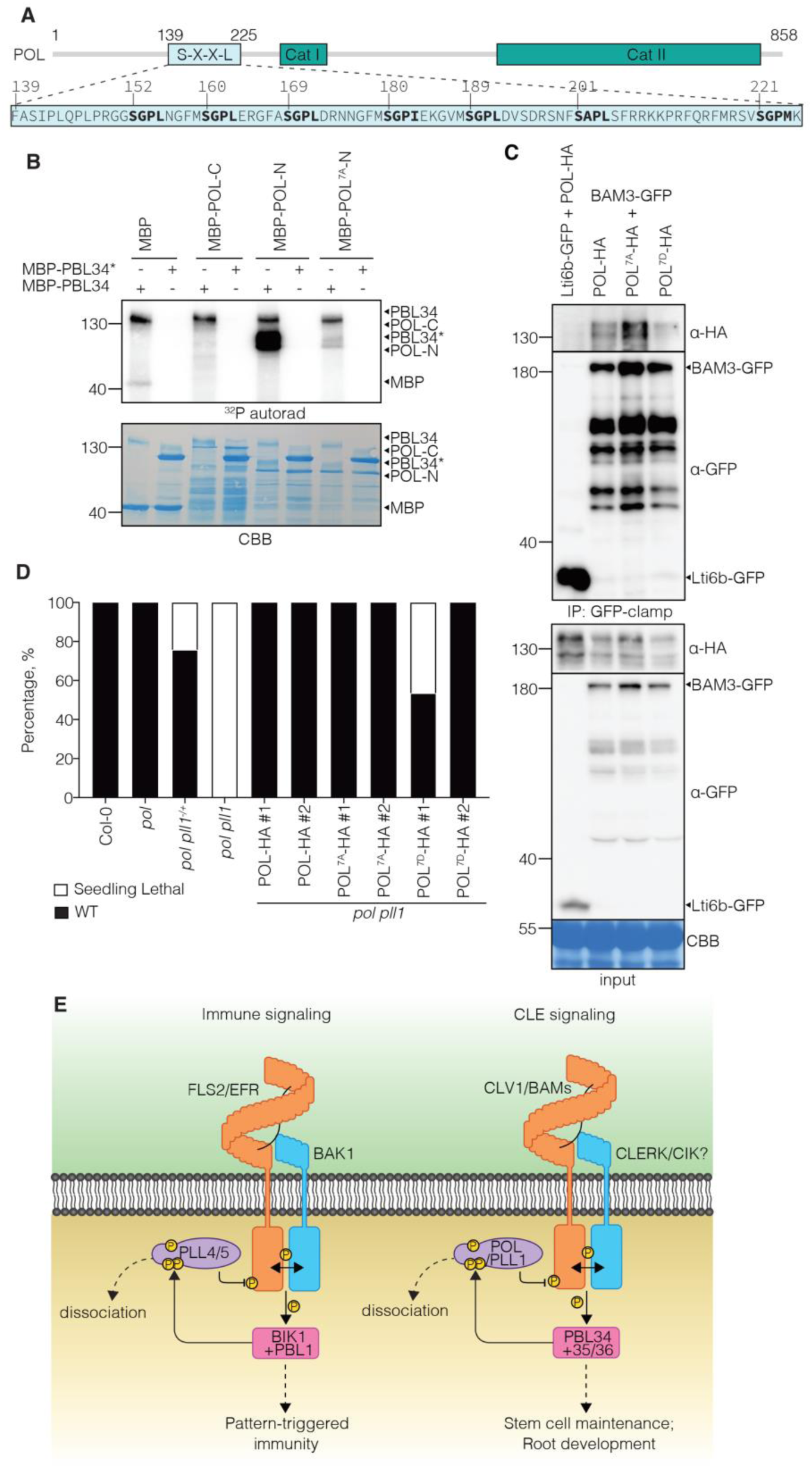
Conservation of the regulatory mechanism in PAMP and CLE receptor complexes. **(A)** Schematic representation of POL domains and details of the S-X-X-L domain of POL. Bold amino acids indicate the S-X-X-L residues targeted for mutagenesis; numbers correspond to the amino acid positions within the protein. **(B)** PBL34 phosphorylates the POL N-terminus in a site-specific manner. Autoradiogram of *in vitro* kinase assay of MBP-tagged C-terminal (C), N-terminal fragments of WT POL (POL-N) or phosphomute variant (POL-N^7A^) with WT PBL34 or inactive (PBL34*). **(C)** BAM3 and POL interact *in planta*. CoIP assay of transiently expressed BAM3-GFP and HA-tagged WT POL, POL^7A^ or POL^7D^ variants in *N. benthamiana* leaves. CBB: Coomassie brilliant blue. **(D)** Quantification of the complementation of *pol pll* phenotype in the shoot by POL^7A^ and POL^7D^ variants. POL-HA and POL^7A^-HA fusion proteins fully complement the seedling lethality of the *pol pll1* double mutant. POL^7D^-HA protein fusion only partially rescues the seedling lethality phenotype. n≥117 plants per genotype. **(E)** Schematic representation of the conserved signaling mechanism between PTI and CLEp signaling pathways exemplified by FLS2 and BAM3 signaling pathways. In the absence of the ligand, PLL family phosphatases damper signaling by inhibiting RK phosphorylation (e.g. PLL4,5 or POL,PLL1). Perception of the apoplastic peptide ligands (e.g. PAMPs or CLEs) by their cognate RKs leads to co-receptor recruitment and activation of specific RLCK-VII isoforms. These RLCK-VII/PBLs phosphorylate PLLs at conserved N-terminal sites, triggering PLL dissociation from the RK complex and appropriate activation of signaling.

To probe the role of POL phosphorylation in CLEp-mediated development, we transformed *pol pll1*^*-/+*^ null mutants with WT or mutant *POL* variants under the native *POL* promoter and isolated viable *pol pll1* doubles to determine ability to complement the seedling lethal phenotype of the double mutant (Song et al., 2006). Consistent with a role for S-X-X-L sites in the phosphorylation-mediated release of POL from primary CLEp receptors, POL and POL^7A^ but not POL^7D^ complemented *pol pll1*, whereas POL^7D^ lines displayed gain-of-function *CLV3*-phenotypes across plant tissues and developmental stages, and in multiple independent transgenic lines (Figure 4D; S7).

Together, these data reveal a conserved regulatory circuit that controls LRR-RK signaling in both immunity and development (Figure 4E), in which specific phosphatases and cytosolic kinases control the activation of ligand-binding receptors in distinct pathways. RLCK-VII isoforms have also been identified downstream of other LRR-RKs in developmental processes, such as SCHENGEN 1 (SGN1)/PBL15, which functions downstream of the receptor GASSHO1(GSO1)/SGN3 to regulate Casparian strip formation (Fujita et al., 2020), further indicating that LRR-RKs share conserved signaling modules downstream of receptor activation. In addition to their roles reported here, RLCK-VII-5 isoforms were also described downstream of LIPOOLIGOSACCHARIDE-SPECIFIC REDUCED ELICITATION (LORE) (Luo et al., 2020), a G-lectin type RK that triggers immunity in response to bacterial 3-OH-FAs (Kutschera et al., 2019). It will be of interest to see if these additional RK pathways may also be regulated by a similar molecular circuitry.

## MATERIALS AND METHODS

### Plant growth and materials

All the mutants investigated in this study are in the *Arabidopsis thaliana* Col-0 wildtype background. Col-0 was used as the control for the phenotypic analyses. The allele named *pbl34-2* in this manuscript carries a C to T point mutation in *PBL34* (At5g15080), which leads to the substitution of leucine 135 by phenylalanine. *pbl34-3* (SALK_126209), *pol-6* (SALK_009469.29.99.f), *pll1-1* (SAIL_319_C08), *clv1-15 (*WiscDsLox489-492B1*), bam1-4 (SALK_107290), bam2-4 (SAIL_1053_E09) pll4* (SALK_203257C), *pll5*(SALK_044162C), *pll4-1 pll5-1* double mutants were obtained from stock centers or described before (Holton et al., 2015). Higher order *rlck-vii* seed stocks were previously described (Rao et al., 2018).

Unless otherwise detailed, Arabidopsis plants were grown in a controlled environment growth chamber at 150 µmol light intensity, 60% relative humidity, and 20°C in a 10-h light cycle. *N. benthamiana* plants were grown in a controlled environment chamber at 120 µmol light intensity, 45-60% relative humidity, and 19-21°C in a 12-h light cycle.

### CLEp-induced root growth inhibition assays

Seeds were sterilized, sown on half-strength Murashige and Skoog media supplemented with 0.3% sucrose and 1% agar, stocked at 4°C for 48h and grown vertically under continuous light at 22°C. Synthetic CLE peptides were obtained from a commercial supplier (Genscript) at > 80% purity, diluted in sterile water and used at the indicated concentration. Root lengths were measured on 600 dpi scans of the plates with Fiji software (Schindelin et al., 2012) using the Simple Neurite Tracer plug-in (Longair et al., 2011).

### CLV3 Peptide Root Elongation Assay

Seeds were sterilized for 10 minutes in 70% ethanol with 0.1% Triton X-100, rinsed in 70% ethanol three times, plated onto ½ Murashige and Skoog (MS-Research Products International), pH 5.7 with 8 g of Phytoagar (RPI) per liter. Seeds were stratified for 48 hours at 4°C. After stratification, seeds were germinated horizontally, under continuous light in a Percival growth chamber set to 22°C for four days. Seedlings at 4 days after germination (DAG) were transferred to vertical ½ MS plates with or without CLV3 peptide (>95% purity, Biomatik) for mock and peptide treatment respectively. Seedlings on vertical plates were allowed to grow for 4 days after transfer, then were scanned and measured using ImageJ software.

### Cloning

PCR products were amplified from plant DNA or plasmid templates (ABRC) using primers listed in Table S1. Mutations were generated using DpnI-mediated site-directed mutagenesis using primers listed in Table S1. For Gateway cloning, PCR products were successively transferred to pDONR vectors by BP reaction (Invitrogen) and pDEST by LR reaction (Invitrogen) according to the manufacturer’s protocols. *pPBL34::gPBL34-CITRINE* was cloned by Gibson strategy (NEB) into a modified pCAMBIA1305,1 plasmid carrying a FASTRED seed selection marker. *POL, PLL1, PLL4*, and *PBL34* fragments were cloned into pOPINM using InFusion (Takara). The *PBL34* promoter was cloned by restriction enzyme cloning into a modified pCAMBIA1305,1 carrying a 3xNLS-VENUS cassette. The *PBL35* and *PBL36* promoters were introduced into a pCAMBIA1305,1 3xNLS-VENUS by Gibson cloning (NEB). The *POL* native promoter (1.6 kb-5’) was integrated into the pMOA34 binary gateway destination vector via PCR and standard restriction cloning.

### Plant transformation

Binary vectors were introduced into Arabidopsis via *Agrobacterium tumefaciens*-mediated (strain GV3101 *pMP90*) transformation by standard floral dipping. Transgenic lines were selected on hygromycin selection media (35 mg/l) or FASTRED seed expression. Single insertion lines were studied.

### *pbl34-2* mutant isolation

*pbl34-2* mutants were isolated as described (Anne et al 2018). CLE26p insensitivity was confirmed in the M3 generation and resistant plants were backcrossed to Col-0. The F1 was uniformly insensitive to CLE26p treatment, suggesting the dominance of this allele. The causative mutation was mapped by whole-genome sequencing of a bulk of 100 seedlings resistant to CLE26p versus 100 sensitive ones as described (Kang and Hardtke, 2016).

### Genotyping

The *pbl34-2* mutation was genotyped with a CAPS strategy. A 730bp PCR product was amplified with the Phire kit (Thermo Fisher). The subsequent PCR product was digested with AflII restriction enzyme, which cuts the wild-type product into 340bp + 390bp fragments but not the *pbl34-2* product. Primers for genotyping are listed in Table S2.

### Root cross sections

7-day-old seedlings were embedded in plastic historesin solution (Technovit 7100). The number of cells per cell layer was quantified at the level of differentiated protophloem, on 10μm cross sections using the Cell Counter plug-in (https://imagej.net/Cell_Counter) of Fiji software (Schindelin et al., 2012). Around 50 roots were used for quantification.

### Protein alignment and phylogenic tree

Protein alignments were performed using CLUSTALW (https://www.ebi.ac.uk/Tools/msa/clustalo/). The output file was uploaded into MEGA X software (https://www.megasoftware.net/) to generate the corresponding phylogenic trees.

### Statistical analyses

Statistical analyses were performed on RStudio software (www.rstudio.com/) or on Prism software (https://www.graphpad.com/scientific-software/prism/). ANOVA analyses followed by Tukey tests were performed with a confidence level of 95%. Specific tests used are indicated in figure captions.

### Recombinant protein expression and purification

All proteins were expressed in *Escherichia coli* strain BL21(DE3) Rosetta pLysS unless otherwise noted. BIK1 or BIK1* (kinase dead, K105A/K106A), PLL4, POL, PLL1, and PBL34 variants expressed as 6xHis-MBP fusion proteins in the pOPINM vector. The EFR cytosolic domain was expressed using pMAL-c4E (MBP-EFR-CD or MBP-EFR*-CD, D849N) in BL21(DE3) Rosetta pLysS or pET-28a(+) (6xHis-EFR-CD) in BL21(DE3)-VR2-pACYC-LamP *E. coli*, respectively. The cytosolic domain of BAM3 WT or kinase dead (BAM3*, D836N) was cloned into a modified pET-28a(+) backbone and expressed as a 6xHis-GST fusion protein (GST-BAM3-CD). All proteins were purified using Amylose Resin (NEB) or HisPur Cobalt Resin (Thermo) for MBP or 6xHis, 6xHis-MBP, and 6xHis-GST fusions, respectively.

### *In vitro* kinase assays

Approximately 1 µg of kinase was incubated with approximately 1 µg of substrate protein in kinase buffer (25 mM Tris-Cl pH 7.4, 5 mM MnCl_2_, 5 mM MgCl_2_, 1 mM DTT). Reactions were initiated by addition of 5 µM ATP plus 0.5 µCi ^32^P-γ-ATP in a final reaction volume of 30 µl. Reactions were carried out at 25 °C for 30 min and stopped by addition of SDS-loading dye and heating at 70 °C for 10 min. Proteins were resolved by SDS-PAGE, transferred to PVDF membrane, and stained with Coomassie brilliant blue G-250. Autoradiographs were imaged using an Amersham Typhoon phosphorimager (GE Healthcare).

For non-radioactive autophosphorylation of EFR, approximately 10 µg of MBP-EFR-CD was incubated in kinase buffer (as above) with 10 µM ATP in a final volume of 100 µl for 1 hour at 25 °C. Free ATP was removed by equilibration into storage buffer (25 mM Tris-Cl pH 7.4, 100 mM NaCl, 10% glycerol, 1 mM DTT) using a 10,000 MWCO centrifugal filtration device (Millipore).

### *In vitro* phosphatase assays

Approximately 1 µg of autophosphorylated MBP-EFR-CD was mixed with approximately 1 µg of phosphatase in buffer (HEPES pH 6.8, 5 mM MgCl_2_, 5 mM MnCl_2_, 200 mM NaCl, 5% glycerol, 1 mM DTT). Reactions were carried out for 90 min at 25 °C. Phosphorylation was monitored by blotting with anti-pThr (anti-phosphothreonine, Cell Signaling Technology 9381, diluted 1:1000 in TBST-5% gelatin from cold water fish skin).

### ROS production assays

ROS burst assays were conducted as previously described (Kadota et al., 2014; Monaghan et al., 2015). For assays with *N. benthamiana*, leaf discs were harvested 24h after infiltration and equilibrated overnight in sterile water and used for assays at 48h after infiltration. For kinetic analyses, RLUs were collected in 30-s intervals for 60 min. T_max_ RLU was defined as the interval with the highest total value.

### Co-immunoprecipitation

Leaves of 4-week-old *Nicotiana benthamiana* leaves were infiltrated with *A. tumefaciens* carrying constructs as indicated in figure captions. In all cases cultures were co-infiltrated with *A. tumefaciens* carrying a P19 suppressor of gene silencing construct. Leaves were detached and bisected 2 days post-infiltration. Leaf halves were equilibrated in liquid MS 1% sucrose (1-2 hours) and subsequently vacuum infiltrated with MS or MS+PAMP as indicated in figure captions. Tissue was frozen and ground in liquid nitrogen. Protein extraction and immunoprecipitation were performed as described previously (Kadota et al., 2014) using GFP-trap (Chromotek) or GFP-clamp (Hansen et al., 2017) resin, as indicated. Proteins were separated by SDS-PAGE and blotted onto PVDF membrane. Membranes were blocked and probed in TBST-5% non-fat milk using anti-GFP (HRP-conjugated B-2, sc-9996 HRP, Santa Cruz, 1:5000 dilution) or anti-HA (HRP-conjugated, 12013819001, Roche, 1:3000 dilution).

### *In vitro* pulldown

Approximately 6 µg each of bait and prey proteins were mixed to 100 µl final volume in buffer (25 mM Tris-Cl pH 7.4, 100 mM NaCl, 0.2% Triton-X, 1 mM DTT). 30 µl was removed (“input”) and the remaining sample was mixed with 50 µl of Amylose Resin (NEB) in a final volume of 500 µl. Samples were mixed at RT for 30 min. The resin was washed three times with buffer and enriched proteins were eluted with 50 µl SDS-loading dye (“pulldown”). Samples were separated by SDS-PAGE, transferred to PVDF, and imaged by blotting with anti-Polyhistidine (Sigma H1029), anti-GST (Upstate 06332), or anti-MBP (NEB E8032) antibodies (all diluted 1:10,000 in TBST-5% non-fat milk powder).

### Confocal Microscopy

5 to 6-day-old seedlings were imaged using an SP8 (Leica) inverted confocal microscope. Samples were prepared in a drop of 0.04 mg/ml propidium iodide solution. CITRINE and VENUS fluorophores were exited at 514 nm and emitted light recorded between 520 and 555 nm. Propidium iodide was excited at 488 and 514 nm and fluorescent light recorded between 600 and 700 nm. CITRINE/VENUS and propidium iodide channels were sequentially acquired. Figures were prepared using Fiji software.

### Immunolocalization

5-day-old-seedlings of transgenic line expressing *pPBL34::gPBL34-GFP* in the *pbl34-3* background were used for whole mount immunolocalization of the PBL34-GFP protein fusion according to (Marhava et al., 2018) and combined with calcofluor white cell wall staining. Primary anti-GFP rabbit (Abcam) antibody was used at 1:600 and secondary Alexa Fluor 546 anti-rabbit antibody was used at 1:500 (Molecular Probes).

### Carpel Counts

Genotype confirmed seeds were sterilized, stratified, and germinated as above. Four DAG, seedlings were transplanted to soil (7 parts top soil to 1 part sand with pesticide) and kept at high humidity for 3-5 days under continuous 24-h light at 23 °C. Seedlings were removed from high humidity and allowed to grow to full maturity, with gentle staking to prevent tangling at ∼3 weeks after transplant. After 5 weeks of growth, the entire number of flowers produced on the primary inflorescence were quantified for carpel number under a dissecting microscope. Data was analyzed in PRISM as above.

### Isolation of *POL* complementation lines

*POL* constructs were introduced into *pol-6 pll1-1/+* line and transgene fixed lines were isolated in the *pol-6 pll1-1* background (or *pol-6 pll1-1/+* if necessary). Fixed lines were used for complementation analysis and expression analysis.

Initial complementation was determined by identifying the ratio of viable to seedling lethal phenotypes for each *POL* variant. The WT and 7A lines displayed 100% viable phenotypes, while the 7D variants displayed a mixed set of phenotypic ratios. Viable 7D lines were transplanted and assayed for post-seedling stem cell defects.

### *POL* Complementation Analysis

For seedling analyses, seeds of *POL* substitution lines were sterilized and plated on ½ MS plates (described above). 8 day old seedlings were assayed for seedling lethal *pol pll* double mutant or WT phenotype. Representative individual plants were genotyped to confirm correct genotype. For adult plant analyses, 5 day-old-seedlings were transplanted to soil (as above). Plants were grown for 4-6 weeks and phenotypes were assayed accordingly.

### Expression Analysis of *POL* Complementation Lines

Bulk 8-day-old seedlings (12 for WT phenotype, 20 for seedling lethal phenotypes, and an approximate weight mix for mixed phenotype) were ground in liquid nitrogen using SPEX Genogrinder and glass beads. The resulting powder was used as the starting material for a standard RNA extraction using a Qiagen Plant Mini Kit (Qiagen). ∼1 ug of RNA from resulting extraction was used as template in standard cDNA synthesis reaction (BioRad iScript cDNA Synthesis Kit). 200 ng of resulting cDNA was used in qRT-PCR reactions to quantify POL expression levels (PowerUp SYBR Green 5x Master Mix-Thermo). CDKA1 was used as an equalization housekeeping gene. Data was analyzed using the ΔΔCT method for three biological replicates with three technical replicates each.

## ACKNOWLEDGEMENTS

The authors thank Jian-Min Zhou (CAS, Beijing) for kindly providing published *rlck-vii* mutants. The Nimchuk lab thanks Tony D. Perdue, director of the University of North Carolina-Chapel Hill Genome Sciences Microscopy Core, for assistance with confocal imaging. The Zipfel group thanks all members for discussions and critical reading of the manuscript.

## Author contributions

Conceptualization: T.A.D., P.A., S.R.J., C.Z., C.S.H, Z.L.N.; Methodology: T.A.D., P.A., S.R.J., C.Z., C.S.H, Z.L.N.; Validation: T.A.D., P.A., S.R.J., A.W.; Formal analysis: T.A.D., P.A., S.R.J., A.W., O.J., Y.G.; Investigation: T.A.D., P.A., S.R.J., A.W., O.J., Y.G., A-M.P., Z.L.N.; Data curation: T.A.D., P.A., S.R.J., A.W.; Writing (original draft): T.A.D., P.A., S.R.J.; Writing (review and editing): T.A.D., P.A., S.R.J., C.Z., C.S.H, Z.L.N.; Visualization: T.A.D., P.A., S.R.J.; Supervision: C.Z., C.S.H., Z.L.N.; Project administration: C.Z., C.S.H., Z.L.N.; Funding acquisition: T.A.D., P.A., C.Z., C.S.H., Z.L.N.

## Data and materials availability

All data is available in the main text or the supplementary materials.

---

**Figure S1.**
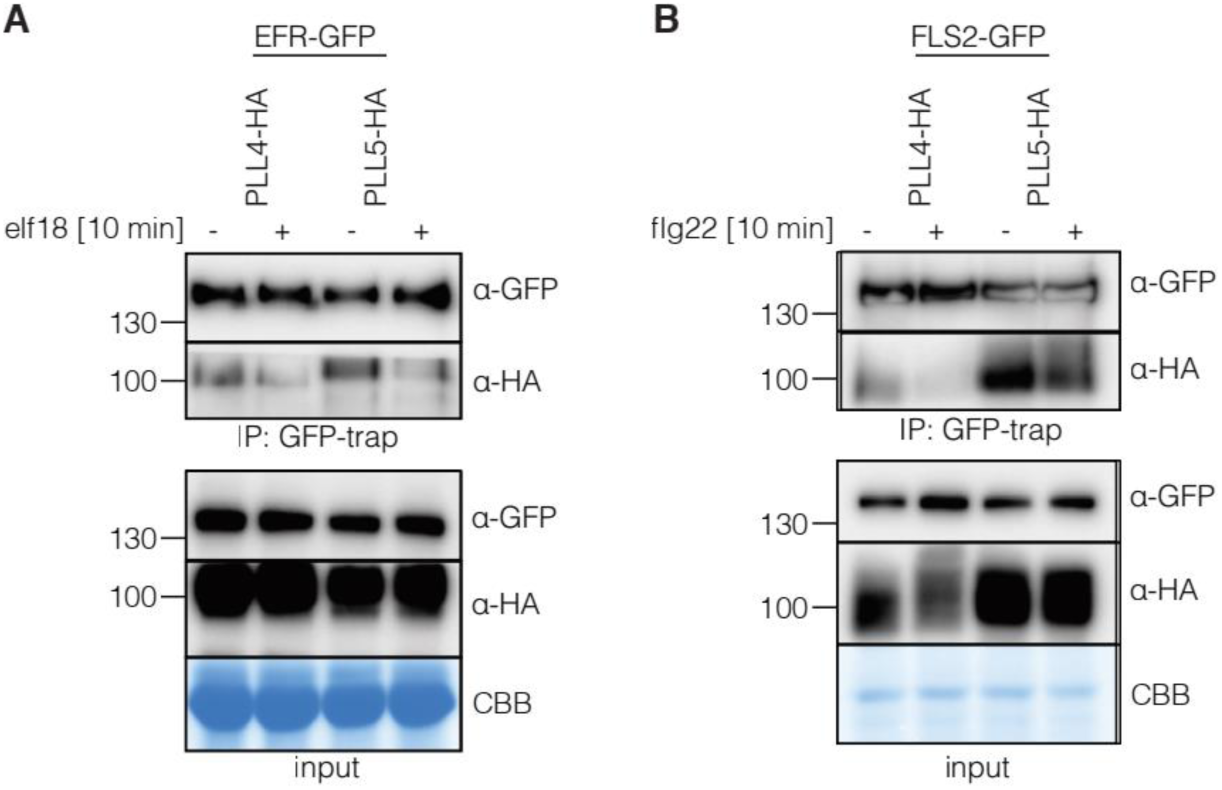
PLL4 and PLL5 dynamically associate with EFR and FLS2 in ligand-dependent manner. **(A)** elf18 triggers EFR/PLL4,5 dissociation *in planta*. CoIP assay of transiently expressed EFR-GFP and HA-tagged PLL4 or PLL5 in *N. benthamiana* leaves with or without treatment with 1 μM of elf18 for 10 minutes. **(B)** flg22 triggers FLS2/PLL4,5 dissociation *in planta*. CoIP assay of transiently expressed FLS2-GFP and HA tagged PLL4 or PLL5 in *N. benthamiana* leaves with or without treatment with 1 μM of flg22 for 10 minutes. CBB: Coomassie brilliant blue.

**Figure S2.**
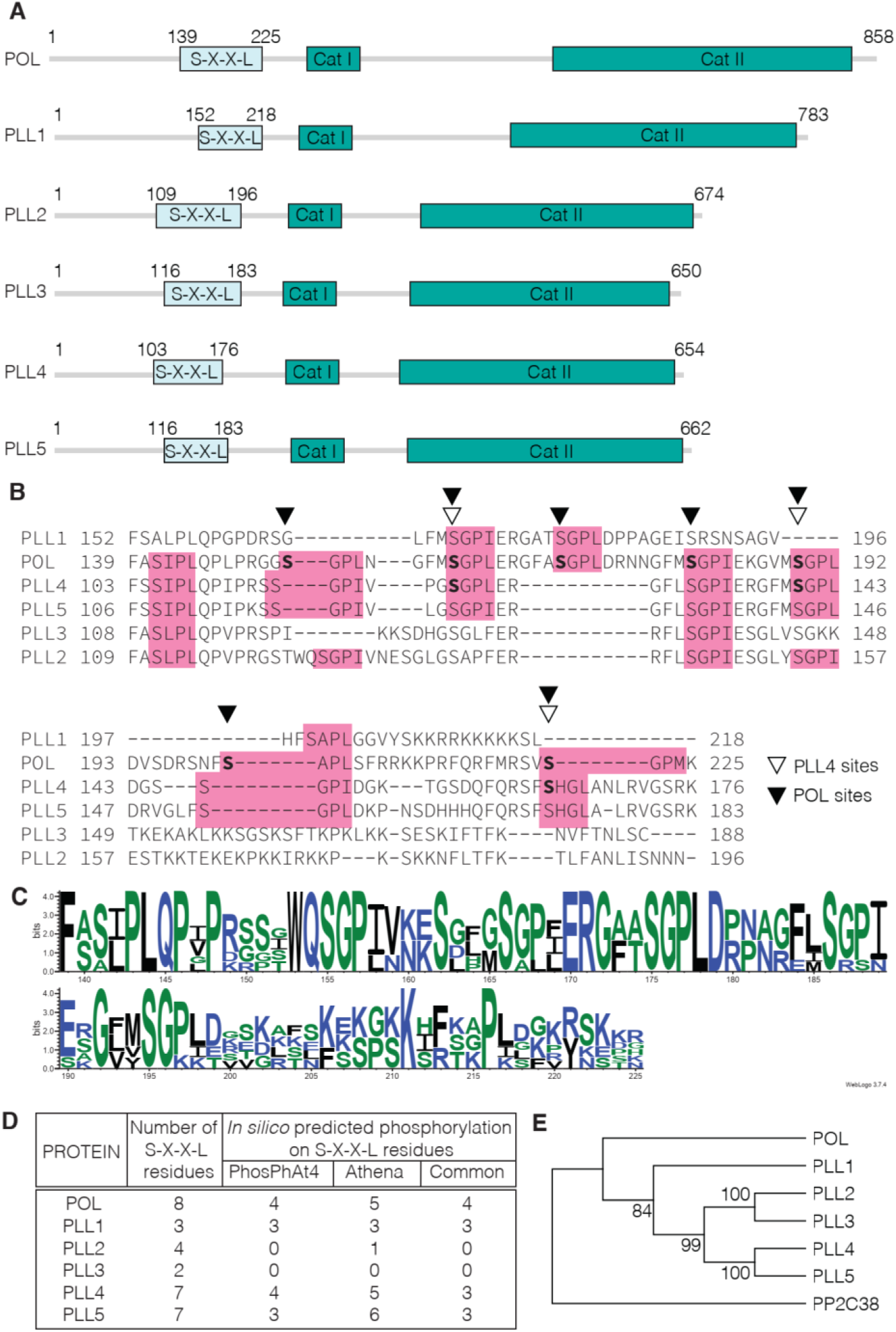
The S-X-X-L domain is conserved among the PLL family. **(A)** Schematic representation of PLLs protein domains. light blue: S-X-X-L domain, green: catalytic domains. Numbers indicates amino acid residues. **(B)** Protein alignment of the S-X-X-L domain of POL family members. Pink square: S-X-X-L residues; bold: residues targeted for mutagenesis in current study in POL (black arrow head) and PLL4 (white arrow head). **(C)** Sequence logo of the S-X-X-L domain of the POL/PLL family created from the alignment in panel B using the WebLogo3 online application (http://weblogo.threeplusone.com). **(D)** Table resuming the *in silico* predicted phosphorylation in the S-X-X-L domain identified in PhosPhAt4 (Heazlewood et al., 2008) and/or Athena databases (Mergner et al., 2020). **(E)** Phylogenetic tree of the POL family based on protein alignment. PP2C38 is used as an outgroup.

**Figure S3.**
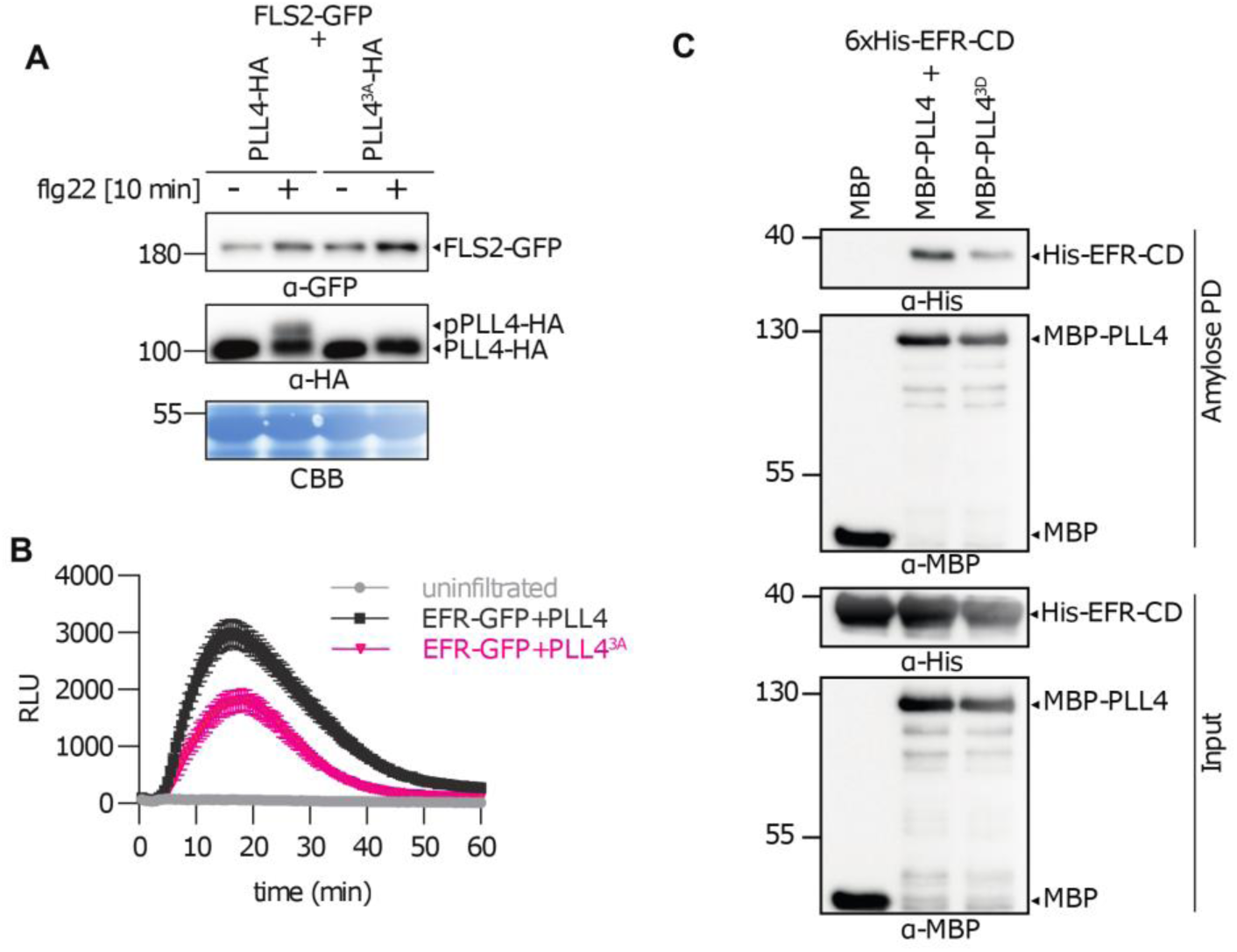
PLL4 regulates PTI in a phosphorylation dependent manner. **(A)** Treatment with 1 µM flg22 induces a phosphosite-dependent mobility shift in PLL4-HA in *N. benthamiana* leaves transiently expressing FLS2-GFP and WT or 3A PLL4-HA. Leaves were treated with or without 1 μM flg22 for 10 min prior to protein extraction and blotting. **(B)** Expression of PLL4^3A^ dampens PTI responses in *N. benthamiana*. ROS burst induction by elf18 (100 nM) on leaf discs of *N. benthamiana* transiently expressing EFR-GFP and WT PLL4 or PLL4^3A^ variants. **(C)** PLL4 phosphomimetic (PLL4^3D^) mutation disrupts direct interaction between PLL4 and EFR-CD *in vitro*. Amylose pulldown assay of 6xHis-tagged cytosolic domain (CD) of EFR with MBP-tagged WT version (PLL4) or phosphovariant (PLL^3D^) of PLL4.

**Figure S4.**
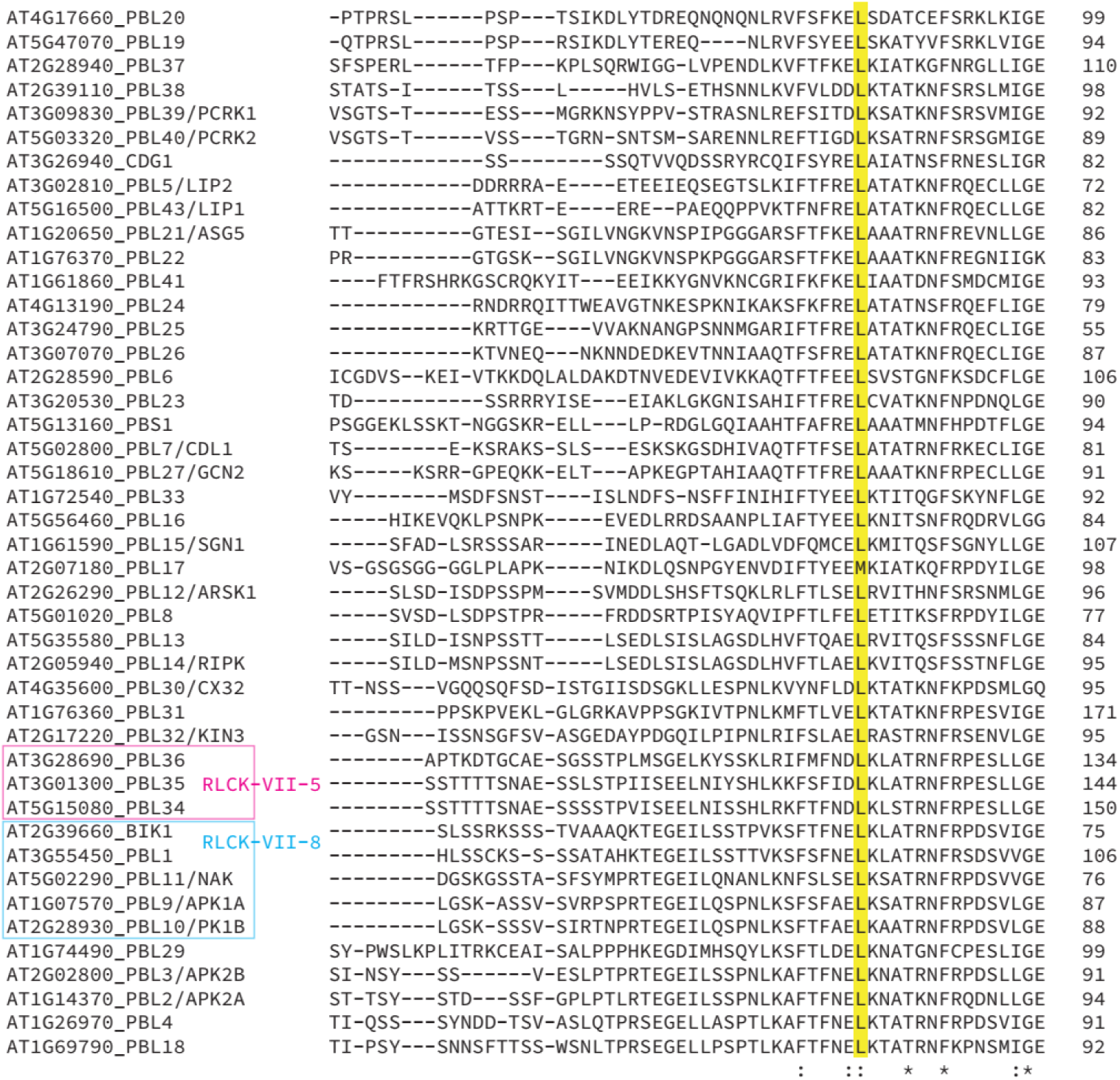
L135F is conserved among the PBL clade. Protein alignment of the PBL family. Magenta square: RLCK-VII-5 clade; cyan square: RLCK-VII-8 clade; yellow square: the highly conserved L residue.

**Fig S5.**
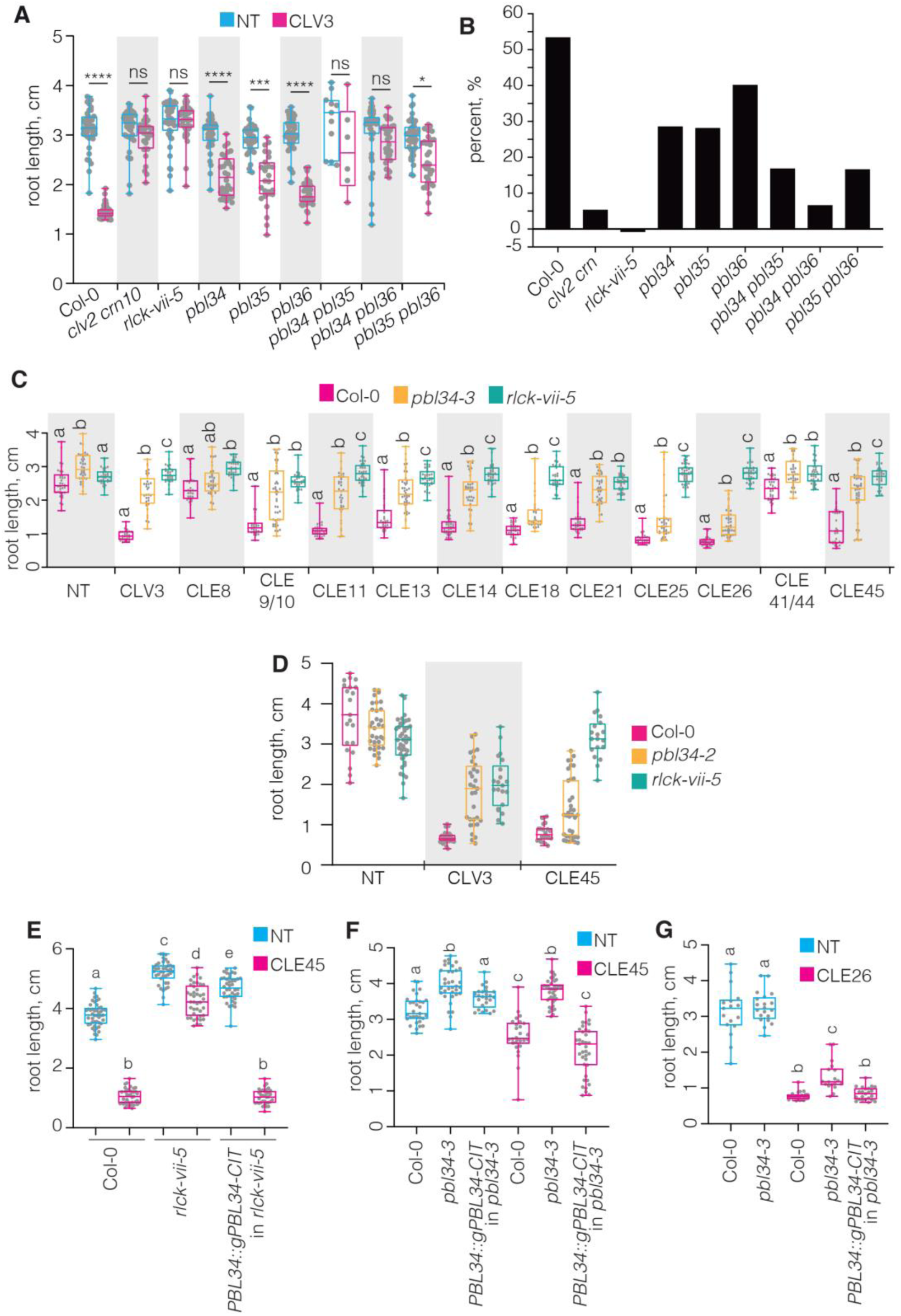
RLCK-VII-5 clade PBL kinases are required for CLE peptide perception. **(A-C)** RLCK-VII-5 members are semi-redundant in the CLEp signaling. (A) Root length of 8-day-old seedlings grown on media with or without 100 nM CLV3p. NT: not treated. Kruskal Wallis Non-Parametric ANOVA test, **** indicates p value <0.0001, * indicates p value <0.001, ns: not significant, n=30. (B) Corresponding growth ratio inhibition calculated as following: 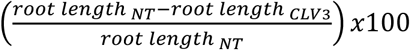. All of the *rlck-vii-5* mutant combinations were less sensitive to CLV3p than the Col-0 background. (C) 7-day-old-seedlings grown on media with 100 nM of indicated CLE peptides. NT: not treated. Letters indicate significant differences within the treatments (ANOVA followed by Tukey test). n= 26-46. **(D)** *rlck-vii-5* is less sensitive to CLV3p and CLE45p treatments than the dominant negative *pbl34-2* mutant. 7-day-old-seedlings grown on media complemented with 100 nM of indicated peptides. NT: not treated. n=19-45. **(E-F)** Complementation of *pbl34-3* mutants expressing *PBL34::gPBL34-GFP* construct. 7-day-old seedlings grown on media complemented with 50 nM CLE peptides. NT: not treated. Letters indicate significant differences within the treatments (ANOVA followed by Tukey test). (E) Complementation assay on CLE26p media. n=17-28. (F). Complementation assay on CLE45p media, n=37-58.

**Figure S6.**
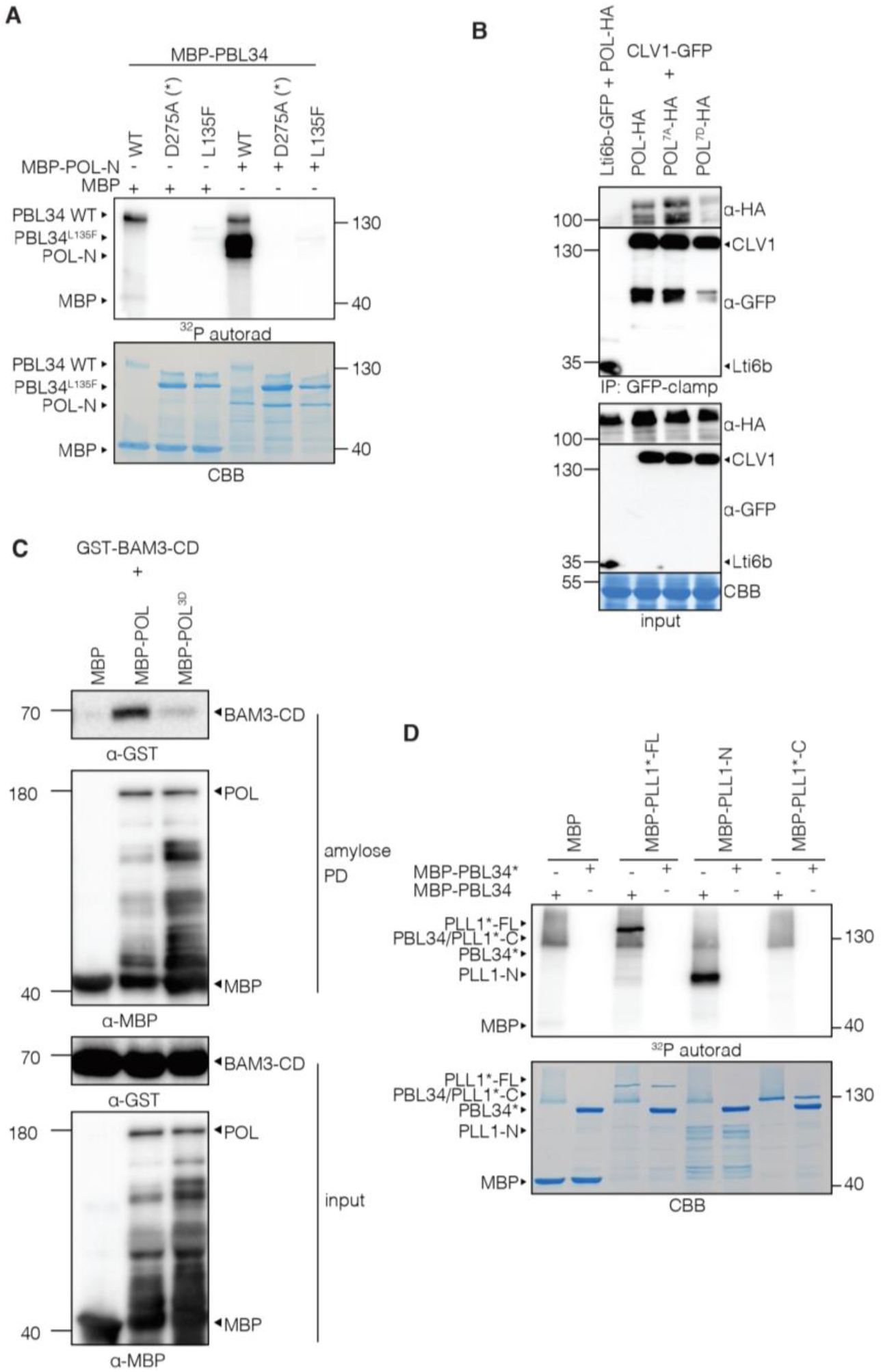
Conservation of RLK-PBL-POL circuitry in CLEp signaling. **(A)** POL is a substrate of active PBL34. Autoradiogram of *in vitro* kinase assay incubating equal amounts of MBP-tagged POL with MBP-tagged WT PBL34 or mutant forms of PBL34 (PBL34^D275A^ or PBL34^L135F^). **(B)** POL phosphorylation status determines its interaction with CLV1 *in planta*. CoIP assay of GFP-tagged CLV1 with HA-tagged WT POL or phosphovariants (POL^7A^ or POL^7D^). (**C**) POL phosphosites control direct interaction with BAM3 *in vitro*. Amylose pulldown assay using equal amounts of GST-tagged cytosolic domain (CD) of BAM3 with MBP-tagged WT (POL) or phosphomimetic (POL^7D^) variants of POL. **(D)** PBL34 phosphorylates PLL1 *in vitro. In vitro* kinase assay incubating equal amounts of MBP-tagged WT version (PBL34) or inactive (PBL34*) of PBL34 recombinant protein with MBP-tagged N-terminus (PLL1-N), catalytically-dead full length (PLL1*-FL), or catalytically-dead C-terminus (PLL1*-C). CBB: Coomassie brilliant blue.

**Figure S7.**
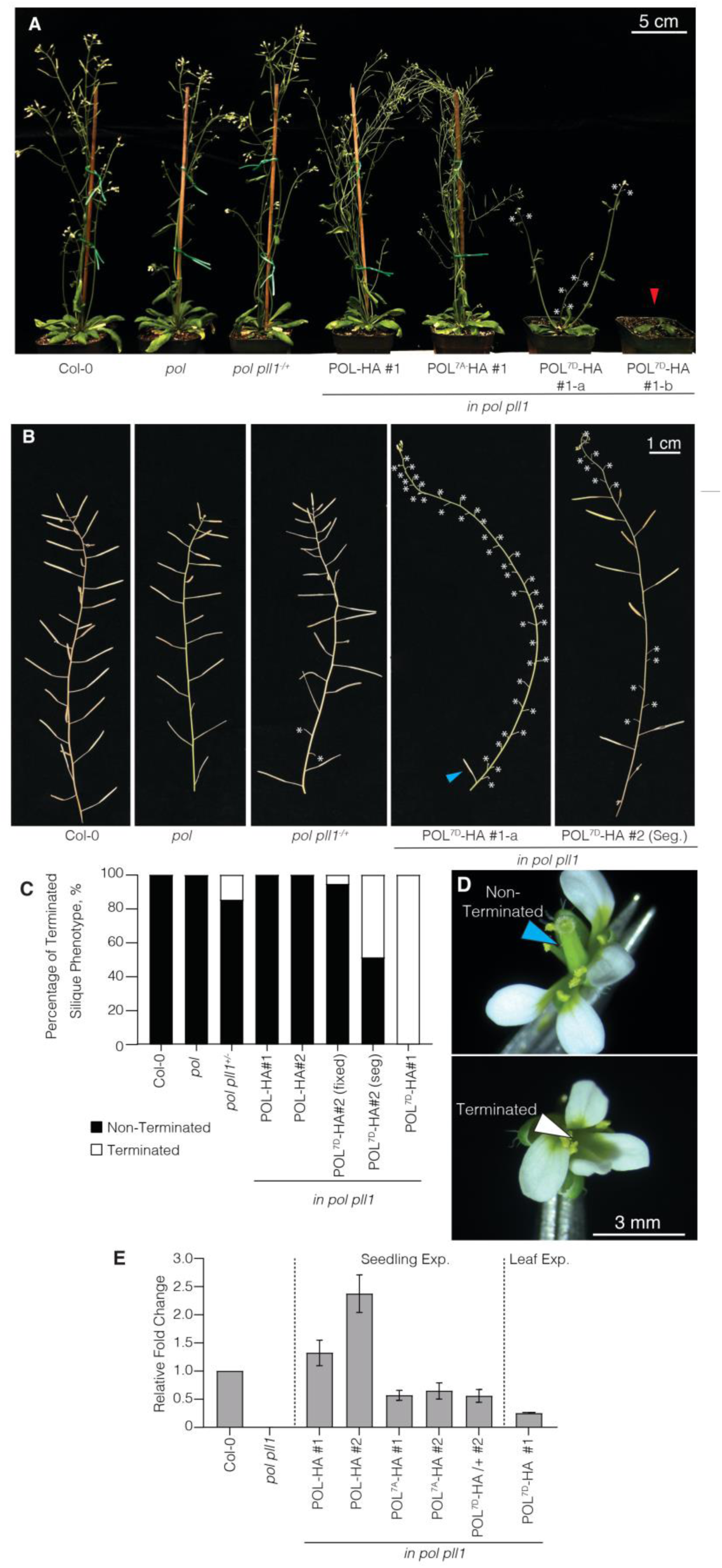
PBL-phosphorylation sites are required for POL function. **(A-C)** POL phosphovariants complement *pol pll1* to varying degrees. (A) Representative pictures of 4-week-old plants expressing different phosphovariants of POL-HA protein fusion (WT, POL^7D^ or POL^7A^) under control of native POL promoter. (B) Representative pictures of 6-week-old stems displaying post-seedling stem cell defect-terminated silique phenotype. white asterisk: terminated silique; cyan arrow head: one successful silique formation for entire line of POL^7D^-HA #1. (C) Corresponding quantification of the shoot complementation based on terminated silique phenotype. STAT. n≥30. **(D)** Detailed pictures of terminated flower compared to WT non-terminated one. cyan arrow head: presence of the pistil; white arrow head: absence of pistil. Scale bar: 3 mm. **(F)** Relative fold change (ΔΔC_T_) expression analysis of *POL* in complementation lines seedlings. Error bars indicate -/+SD.

**Table S1.**
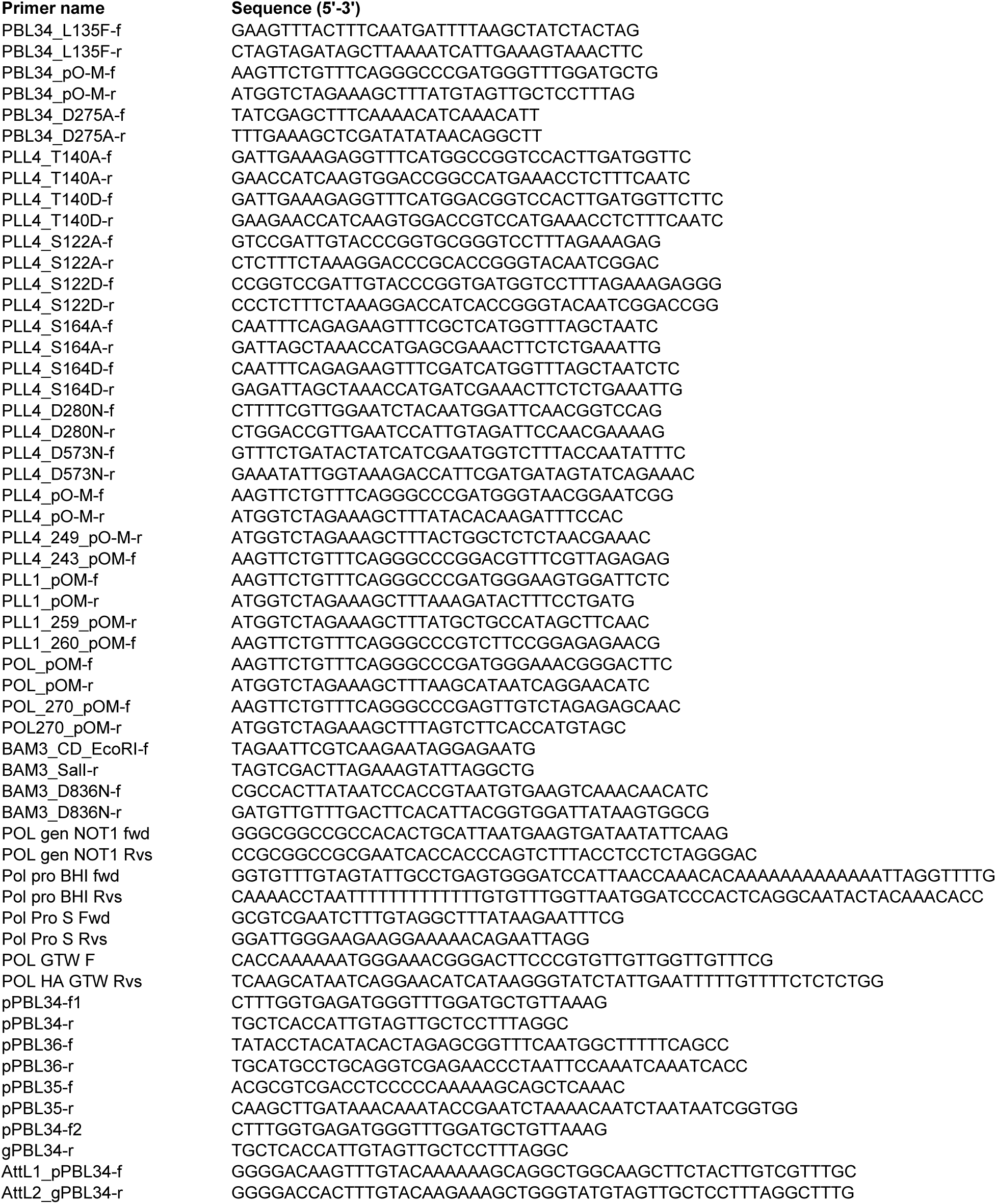
Primers used for cloning in this study.

**Table S2.**
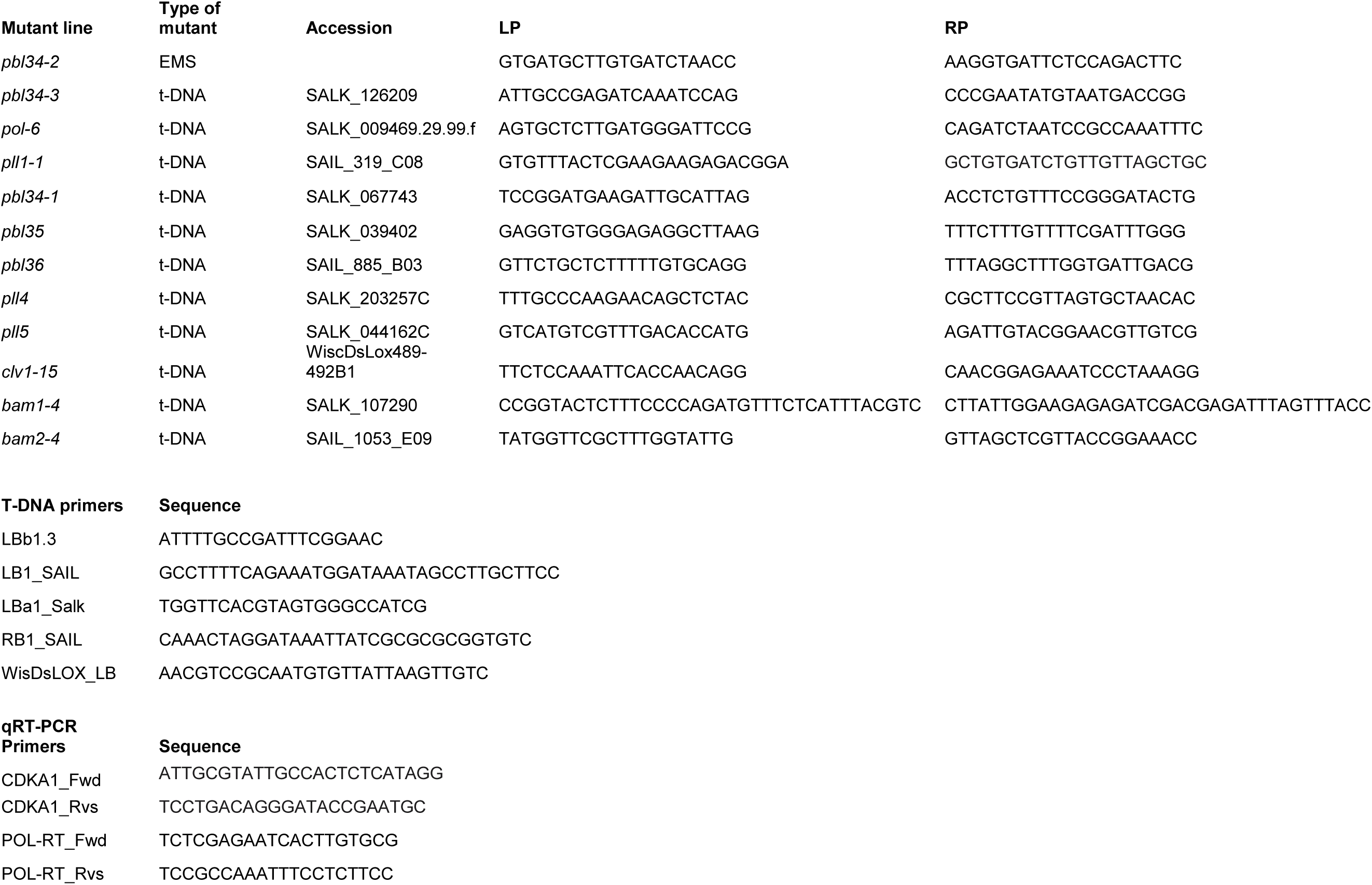
Primers used for genotyping and qPCR in this study.

